# Intrinsic dimensionality of deep learning representations reveals cell death-associated heterogeneity in Parkinson’s disease iPSC-derived neurons

**DOI:** 10.64898/2026.10.02.756370

**Authors:** Seohyeon Kang, Jisung Park, Minee L. Choi

## Abstract

As AI is increasingly used to stratify heterogeneous Parkinson’s disease, it is essential to determine whether learned representations preserve disease-relevant variation within diagnostic or genetic groups. In iPSC-derived neurons from three familial PD cases, deep learning representations encoded an independent measure of cell death, and their intrinsic dimensionality increased with death burden. Latent dimensionality offers a way to examine heterogeneity before AI-derived clusters are interpreted as disease subtypes.

## Main

Parkinson’s disease (PD) is heterogeneous not only in clinical course but in the cellular processes that drive degeneration^1^. Mutations including SNCA triplication (SNCAX3), PINK1 loss of function and GBA variants converge on partially overlapping mitochondrial and lysosomal phenotypes^2–4^, and this overlap makes subtype-associated cellular states difficult to resolve by conventional morphological measurement. Patient-derived induced pluripotent stem cell (iPSC) models preserve disease-relevant genetic backgrounds while permitting controlled high-content imaging, and deep learning applied to such images can predict mechanistic subtypes of PD in patient-derived neurons with high accuracy^5^, distinguish PD genotypes reproducibly across independent laboratories^6^, and recover organelle-level phenotypes more generally^7^.

Deep learning is increasingly used to stratify heterogeneous patient populations, and clusters recovered from learned representations are readily interpreted as candidate disease subtypes. Image-based networks classify mechanistic PD subtypes at accuracies approaching 95%^5^. Previous studies have sought to interpret the biological information encoded by learned representations, but the structure of the representation itself and disease-relevant variation retained within labelled groups remain less explored. We therefore asked whether the geometry of the learned representation could quantify this within-group variation. Under the manifold hypothesis, high-dimensional data lie near lower-dimensional manifolds describable by a limited number of effective degrees of freedom^8^. The intrinsic dimensionality (ID) of a representation estimates that number, and has been used to characterise image representations and transcriptional state during differentiation^9,10^. If degenerating neurons occupy progressively more varied cellular states, ID should increase with degeneration; if degeneration follows a single stereotyped trajectory, it should not. Here we test this in convolutional neural network (CNN) representations of live-cell images of iPSC-derived cortical neurons from healthy controls and three PD genetic subtypes.

We used a previously reported live-cell imaging dataset of cortical neurons differentiated from three control and three PD iPSC lines carrying SNCAX3, GBA or PINK1 mutations^11^. Nuclei, mitochondria and lysosomes were labelled with Hoechst 33342, TMRM and LysoTracker; SYTOX Green marked dead cells and was withheld from all CNN training (Fig. 1a). Latent representations were generated using two encoders, ResNet18 and SupMoCo; the latter incorporated supervised contrastive learning with a momentum-updated key encoder^11^ (Fig. 1b). Both separated the four groups with comparable accuracy (approximately 89% for ResNet18 and 90% for SupMoCo; Fig. 1c), and UMAP projections showed separation of controls from the three subtypes (Fig. 1d). This is consistent with recent work showing that image-based machine learning separates PD genotypes in iPSC-derived neurons reproducibly across independent laboratories^6^.

**Fig. 1.**
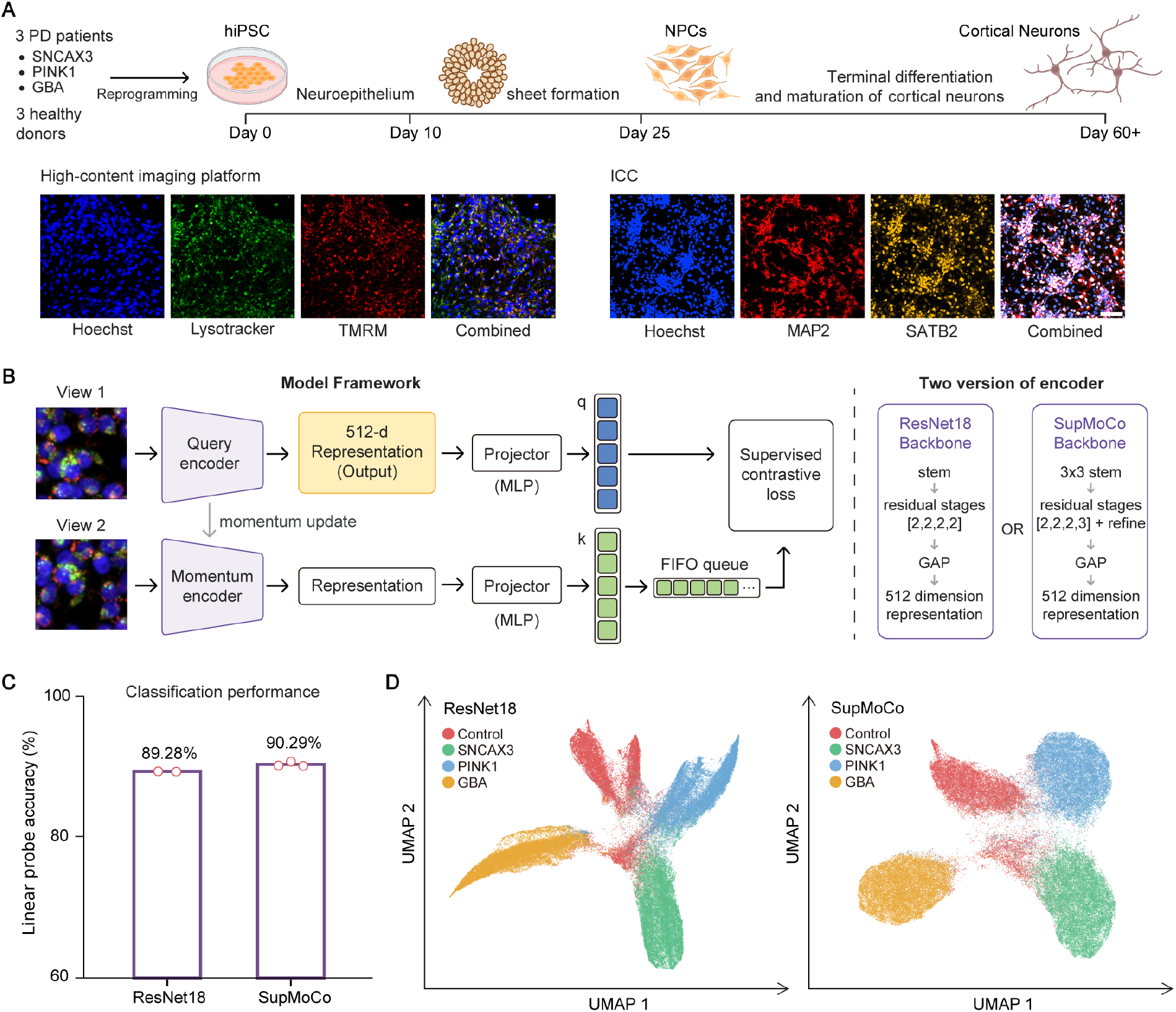
CNN representations distinguish Parkinson’s disease subtypes from live-cell neuronal images. a, Cortical neuron differentiation and imaging workflow using hiPSCs from PD patients and healthy donors. Cortical neuron identity was confirmed by immunocytochemistry for MAP2 and SATB2; Hoechst, LysoTracker and TMRM were used for live-cell imaging. b, Representation learning by supervised momentum contrastive learning. Two augmented views were passed to the query and momentum encoders. Two architectures were used, ResNet18 and a custom SupMoCo backbone, and 512-dimensional encoder outputs taken before the projection head were used for downstream analysis. c, Linear-probe classification accuracy for each encoder; each point is one model trained with a different random seed. d, UMAP visualisation of ResNet18 and SupMoCo representations; each point is one image patch.

We next asked whether cell-death burden, which was withheld during training, could be recovered from the representations. Ridge regression was performed independently within GBA, PINK1 and SNCAX3, giving mean held-out R^2^ values of 0.300 for ResNet18 and 0.533 for SupMoCo (Fig. 2a). Performance varied by subtype and was lowest for GBA (Fig. 2b), consistent with GBA-associated alterations being coupled to terminal cell death less directly than those in PINK1 or SNCAX3. Because regression was performed within genotype, this predictive performance cannot be attributed to between-genotype differences. Cellular morphology of dying cells is partially visible in the input channels, so some of this signal is expected; the relevant observation is that it is retained in representations trained only to separate genotypes. Overlaying SYTOX burden post hoc on UMAP embeddings revealed a graded distribution of cell-death burden within genotype-associated regions (Fig. 2c). PHATE embeddings further revealed a trajectory-like organisation of cell-death burden across latent states (Fig. 2d), consistent with its ability to preserve continuous biological trajectories in high-dimensional data^12^.

**Fig. 2.**
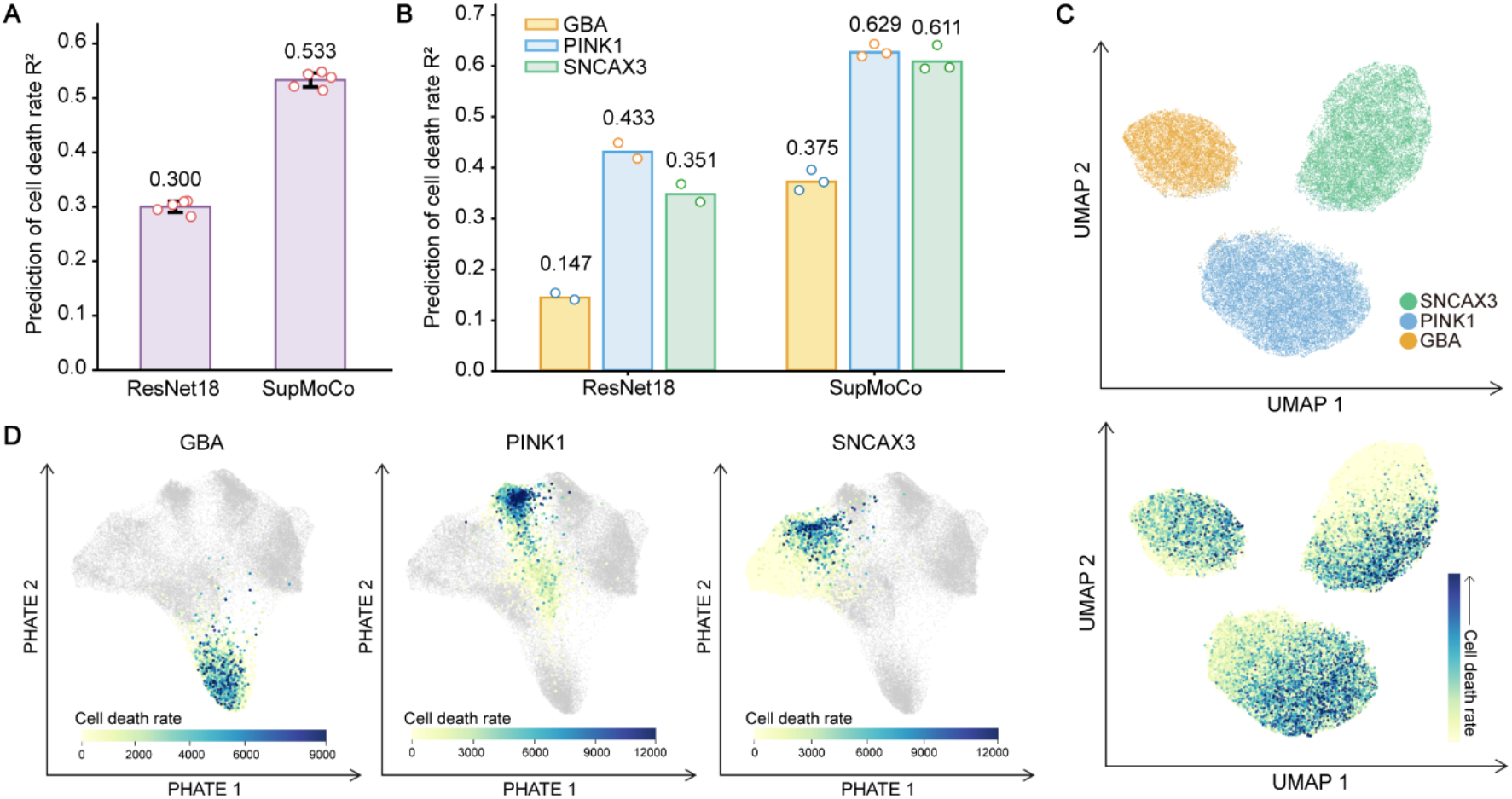
Cell-death burden is encoded in CNN latent representations. a, SYTOX-based cell-death burden predicted by Ridge regression from ResNet18 and SupMoCo representations. Each dot is the mean held-out test R^2^ from one repeated train/test split, averaged across encoder seeds and PD genotypes; bars indicate the mean across five repeated splits. b, Ridge regression performed separately for GBA, PINK1 and SNCAX3; each point is one model trained with a different random seed. c, UMAP visualisation of SupMoCo representations for the three PD subtypes. Upper panels are coloured by genotype; lower panels show SYTOX-based cell-death burden overlaid post hoc on the same coordinates. d, Shared PHATE embedding generated from all control, GBA, PINK1 and SNCAX3 samples. Cell-death burden is shown separately for each PD subtype; grey points are representations from the other subtypes.

We then quantified the effective degrees of freedom underlying this variation. Latent representations were sorted by SYTOX burden within each subtype and grouped into overlapping windows, and ID was estimated per window. Representational structure was reproducible across independently trained models, as assessed by centred kernel alignment (Fig. 3a) and by correlation of pairwise latent distances between the same image tiles (Fig. 3b), indicating that the geometry examined below is not specific to a single random initialisation.

**Fig. 3.**
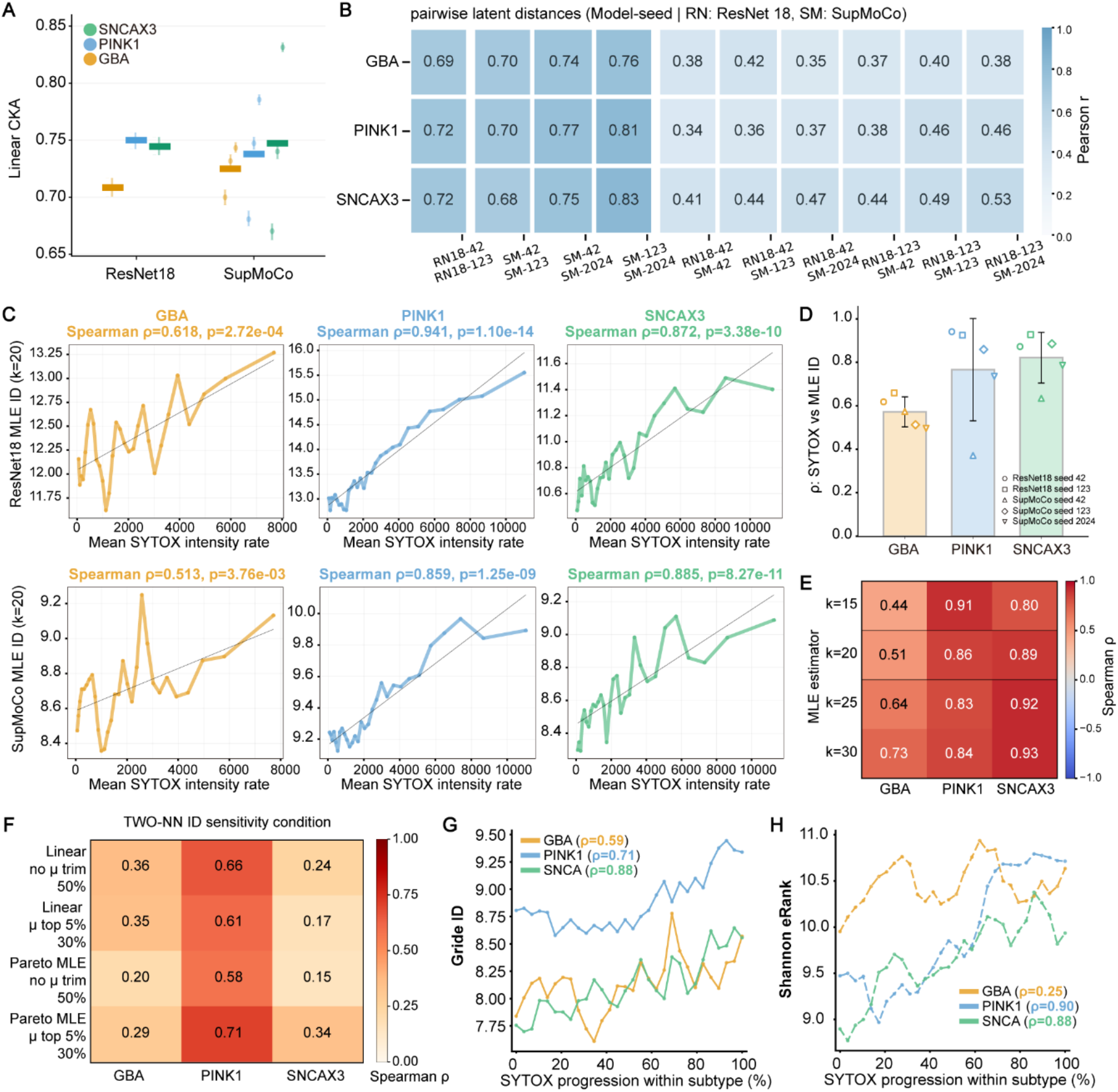
Intrinsic dimensionality increases with cell-death burden across models and estimators. a, Linear CKA between representations from different encoder architectures and random seeds; points indicate similarity for individual seeds. b, Pairwise distances between identical image patch pairs computed in each model’s latent space, compared across seeds by Pearson correlation. c, ID versus SYTOX burden using the Levina–Bickel MLE estimator (k = 20); upper panels, ResNet18; lower panels, SupMoCo. Each point is a window-level ID estimate from a SYTOX-ranked overlapping window. Spearman ρ and p-value are shown per panel. d, Spearman correlations between SYTOX burden and MLE ID across architectures and seeds; bars indicate the subtype mean and symbols individual architecture-seed combinations. e–h, Alternative dimensionality analyses performed using SupMoCo representations. e, Sensitivity of the ID–SYTOX association to the MLE neighbourhood parameter (k = 15, 20, 25, 30). f, Sensitivity of the association under different fitting and trimming conditions for TWO-NN. g, ID along SYTOX progression estimated by GRIDE. h, Shannon effective rank along SYTOX progression.

Using the Levina-Bickel maximum likelihood estimator (MLE)^13^, ID increased with higher SYTOX burden in both ResNet18 and SupMoCo representations (Fig. 3c,d), and the association was preserved across different numbers of nearest neighbours (k) and different window size. (Fig. 3e, Supplementary Fig 1.). To test whether this relationship depended on the estimator, we applied other complementary approaches. TWO-NN^14,15^ reproduced the positive association with greater variability across conditions, reproduced the positive association with greater variability across conditions, which may arise from its dependence on the ratio of the two nearest-neighbour distances and sensitivity to local sampling fluctuations (Fig. 3f). GRIDE^15^, which extends the ratio framework to a broader neighbourhood scale, again showed increasing ID (Fig. 3g). Finally, Shannon effective rank^16^, which measures how broadly variance is distributed across feature directions from the latent covariance spectrum rather than from local neighbourhoods, also increased with cell-death burden (Fig. 3h). The association therefore holds across estimators with different underlying assumptions.

Complementary geometric measures indicated that this increase does not reflect simple expansion along one axis (Supplementary Fig. 2). The median 10-nearest-neighbour radius decreased with cell-death burden in all three subtypes, and the orientation difference between neighbouring local tangent subspaces increased, whereas Laplace–Beltrami mean curvature showed no consistent change. Higher cell-death burden was thus associated with locally denser but less consistently oriented representation structure, a pattern consistent with variation organised into multiple directions rather than along a single severity axis.

Together these results indicate that CNN representations of PD iPSC-derived neurons retain cell-death-associated information that was never supplied during training, and that the dimensional complexity of those representations rises as cell-death burden rises. One interpretation is biological: cell death proceeds through diverse and interacting pathways^17,18^, and combinations of damage processes may present as varied organelle-level states, so that neurons summarised by a single cell-death value nonetheless occupy materially different cellular states. On this reading ID supplies a quantitative index of state diversity that a scalar death metric cannot capture (Fig. 4). Geometric analysis of phenotypic image representations offers a quantitative cellular heterogeneity without cell-death labels in patient-derived disease models, complementing classification-based approaches that presuppose the categories of interest. Whereas image-based classifiers separate PD genotypes and identify the organelle features that drive that separation^6^, the geometry of the same representations describes variation that persists within a genotype, which classification accuracy does not report. As AI-based stratification of Parkinson’s disease expands, latent dimensionality provides a practical way to examine whether disease-relevant heterogeneity is preserved within a group before clusters recovered from such representations are interpreted as candidate disease subtypes.

**Fig. 4.**
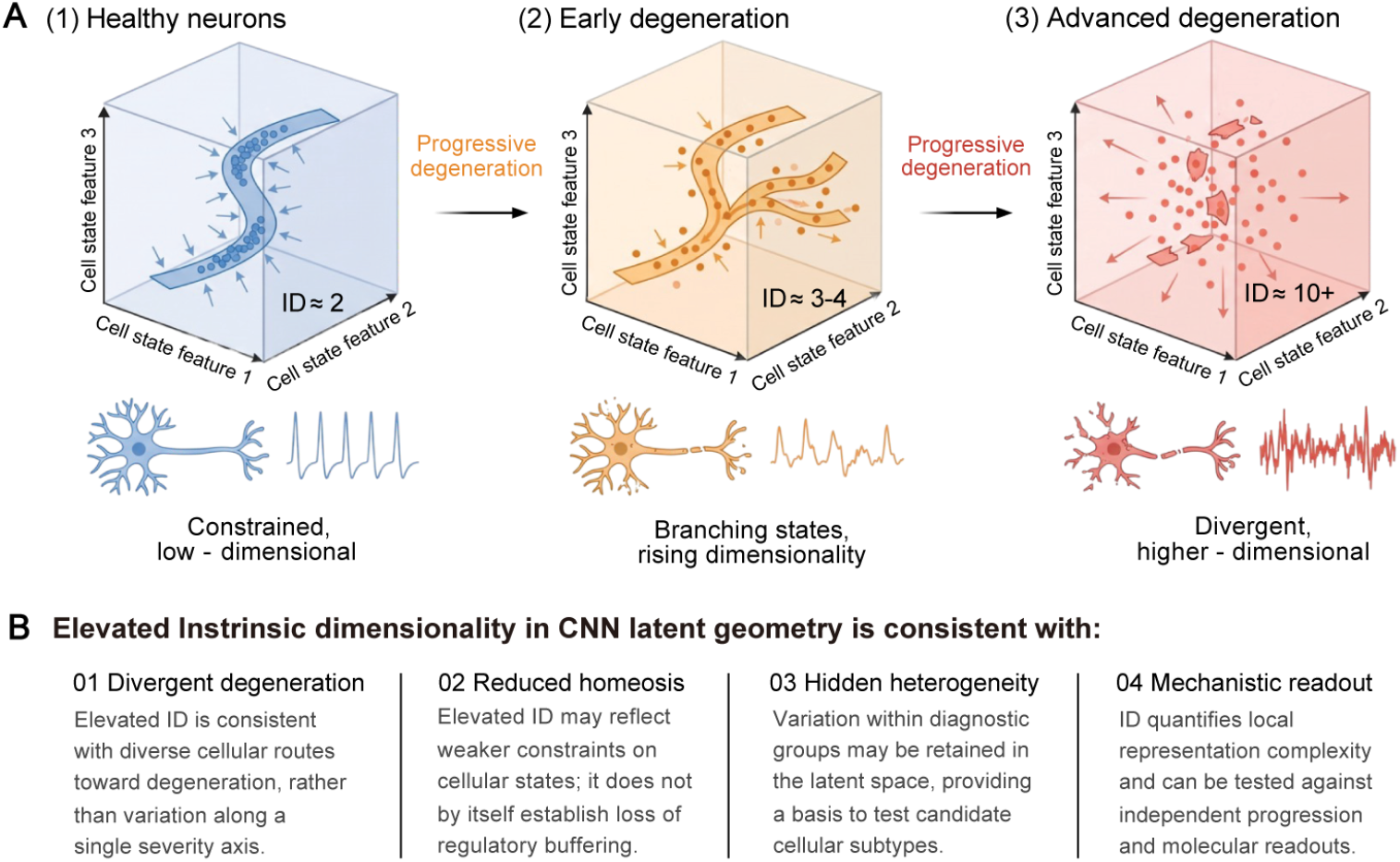
Interpretation of elevated intrinsic dimensionality in CNN latent geometry. a, Schematic of how latent cell-state geometry is proposed to change with progressive degeneration. Each cube represents a latent space spanned by cell-state features, points represent individual image representations, and arrows indicate constraints on accessible states. States are tightly constrained in healthy neurons (ID ≈ 2), branch into multiple trajectories in early degeneration (ID ≈ 3–4), and disperse across many directions in advanced degeneration (ID ≈ 10+). ID values are illustrative and indicate relative differences rather than measured estimates. b, Four interpretations consistent with the increase in intrinsic dimensionality (ID) observed with rising cell-death burden: divergent routes toward degeneration rather than a single severity axis, weaker constraints on cellular states, heterogeneity retained within diagnostic groups, and ID as a testable readout of local representation complexity.

This study has several limitations. Although we interpret increased intrinsic dimensionality as reflecting biological state complexity, CNN latent representations are shaped by multiple image features and biological factors during learning, and geometric properties of representations may be influenced not only by the structure of the data but also by model architecture. The consistency of the changes with cell-death burden, an independent biological measurement that was not used during training, and across two architectures, multiple random seeds and four estimators supports a biological rather than a purely architectural origin, but does not establish one. Intrinsic dimensionality estimators are in addition sensitive to sample density, image noise and the compositional heterogeneity of the data to which they are applied, and images of degenerating neurons are both noisier and drawn from a more variable population than images of healthy neurons; label-permutation and density-matched control analyses will be needed to separate biological state diversity from measurement variance. A further constraint is that each genetic subtype here derives from a single donor line, so subtype-level comparisons are confounded with donor background. We restricted all quantitative analyses to within-genotype comparisons for this reason, but replication across additional lines, ideally isogenic pairs, is required, at the scale of multi-line multi-site designs recently applied to this problem. Future studies connecting interpretable image features, such as mitochondrial, lysosomal and nuclear morphology, or transcriptomic and functional profiles, with latent dimensionality would further clarify the biological basis of increased representation complexity.

## Methods

### Image dataset

We used a previously published image dataset^11^ comprising three control and three Parkinson’s disease patient-derived iPSC lines. While the original dataset included SNCA triplication, GBA, and LRRK2 patient lines, we selected SNCA triplication, GBA, and PINK1 patient lines for the present study (Supplementary Table 1). All lines were differentiated into cortical neurons using a modified version of the protocol of Shi et al.^19^ Briefly, when iPSCs grown on Matrigel-coated 6-well plates reached 90–100% confluence, neural induction was initiated in N2B27 medium supplemented with 10 μM SB431542 and 1 μM dorsomorphin, refreshed daily for 10 days. Neuroepithelial sheets forming between days 9 and 10 were split with Dispase and replated in neural induction medium. Neuronal identity was confirmed by MAP2 and SATB2 immunocytochemistry. For live-cell imaging, nuclei, mitochondria, lysosomes and dead cells were labelled with Hoechst 33342, TMRM, LysoTracker and SYTOX Green respectively.

### Image preprocessing and CNN representation learning

Live-cell images were analysed after maximum intensity projection and divided into non-overlapping 128 × 128-pixel patches treated as independent inputs. Quality filtering removed patches with low signal, excessive background, saturation or blur. Two encoders were used: a standard ResNet18 backbone with the final classification layer removed, using the global-average-pooled output as the representation; and a custom residual encoder in the SupMoCo framework^11^. Only the Hoechst, TMRM and LysoTracker channels were used as input; the SYTOX channel was never seen by either encoder. Both models were trained from random initialisation with SGD (initial learning rate 0.1, momentum 0.995, temperature 0.07, weight decay 1 × 10^−4^, batch size 512, up to 100 epochs), with a 4-epoch linear warm-up followed by a cosine schedule. Augmentation comprised random 90° rotations and horizontal/vertical flipping for ResNet18, and random 90° rotations for SupMoCo; channel-wise instance normalisation was applied for both. L2 normalisation was applied to contrastive embeddings during training, whereas downstream analyses used raw 512-dimensional encoder outputs taken before the projection head. ResNet18 was trained with two random seeds and SupMoCo with three.

### Cell-death quantification and prediction

Cell death was quantified per image patch using SYTOX Green fluorescence. Nuclei were segmented from the Hoechst channel with a pretrained StarDist2D model and matched to the corresponding SYTOX image from the same tile. Edge patches were excluded, and only patches in which the nucleus mask occupied a minimum fraction of the patch area were retained. Ridge regression was performed independently for GBA, PINK1 and SNCAX3. Group-aware splitting ensured that tiles from the same source image or acquisition group did not appear in both training and test sets, and performance is reported as held-out R^2^ averaged over five repeated splits and across encoder seeds.

### Intrinsic dimensionality analysis

For each subtype, samples were sorted in ascending order of SYTOX value and overlapping windows corresponding to 10% of the total sample number were constructed, giving 30 windows per subtype. ID was estimated within each window. The Levina– Bickel maximum likelihood estimator^13^ was the primary estimator, with k = 20 for the main analysis and k = 15, 25 and 30 for sensitivity analysis; window ID was the mean of valid local estimates. TWO-NN^15^ was computed from the ratio of first- and second-nearest-neighbour distances, and GRIDE^15^ from a generalised ratio over a broader neighbourhood range. Shannon effective rank^16^ was computed from the normalised eigenvalue spectrum of the mean-centred latent covariance matrix within each window. Representational reproducibility across seeds was assessed by linear centred kernel alignment and by Pearson correlation of pairwise latent distances computed on identical image-patch pairs. Formal definitions of all estimators are given in Supplementary Methods.

### Exploratory latent geometry

Within the same SYTOX-ranked windows, we computed the median Euclidean distance from each point to its 10th nearest neighbour; Laplace–Beltrami mean curvature magnitude from a k-nearest-neighbour graph with an adaptive Gaussian kernel and Coifman–Lafon normalisation; and local tangent-subspace orientation change, obtained by reducing representations to 50 dimensions by global PCA, applying local PCA over the 30 nearest neighbours of each point, and quantifying orientation differences between neighbouring subspaces by chordal distance^20^. Monotonic associations with SYTOX burden were evaluated by Spearman’s ρ. Details are given in Supplementary Methods.

### Statistics and reproducibility

Associations between window-level geometric measures and SYTOX burden were quantified by Spearman’s rank correlation coefficient. These analyses were considered exploratory; therefore, correlation coefficients were used as the primary measure of association, while p-values are provided as descriptive statistics without multiple-comparison correction.

## Data availability

The imaging dataset analysed in this study was partially derived from a previously described dataset^11^, with the PINK1 mutant line additionally included in this study. The full dataset has been deposited in Zenodo and will be made publicly available upon publication of the manuscript.

## Code availability

All analysis code is available at https://github.com/Seohyeon-1126/PD-neuron-intrinsic-dimensionality.git and archived at 10.5281/zenodo.23004622

## Acknowledgements

This work was supported by the National Research Foundation of Korea, funded by the Korean government (RS-2023-00266872, RS-2024-00343012 to M.L.C.), and by the InnoCORE program of the Ministry of Science and ICT (N10260004, N10260159). M.L.C. acknowledges support from the KAIST–Formosa R&D Center.

## Author contributions

S.K., J.P. and M.L.C. conceived and designed the study. S.K. developed the analysis code, performed the analyses and wrote the original draft. J.P. contributed to conceptualization, provided critical input and edited the manuscript. M.L.C. supervised the study and wrote and edited the manuscript. All authors reviewed and approved the final manuscript.

