## Supplementary data for "Intrinsic dimensionality of deep learning representations reveals cell death-associated heterogeneity in Parkinson’s disease iPSC-derived neurons"

| Line ID | Mutation | Diagnosis | Sex / Age | Source | Independent differentiations (n) | QC-passed cropped images (n) |
|---|---|---|---|---|---|---|
| **C4-NTN** | N/A | Control | Male / 51 | Cedars Sinai | 2 | 34,160 |
| **WTSli018-A** | N/A | Control | Male / 55-59 | EBiSC | 3 | 51,378 |
| **WTSli019-B** | N/A | Control | Male / 55-59 | EBiSC | 6 | 90,659 |
| **AST18** | SNCA triplication | PD | Female / 55 | UCL (Tilo's lab) | 3 | 44,121 |
| **ND50093** | PINK1: P.I368N/P.I368N | PD | Female / 60 | NINDS | 4 | 80,598 |
| **N370S/WT** | GBA N370S/WT | PD | Female / 53-55 | UCL (Schapira lab) | 3 | 68,522 |

**Table S1. Characteristics of the hiPSC lines and imaging dataset used in this study.** Summary of the hiPSC lines, genetic background, donor information, cell-line source, number of independent differentiations, and image patches used for downstream analysis. Imaging data from each cell line were collected across multiple independent differentiations, collectively yielding more than 300,000 image patches for representation learning and downstream analyses.

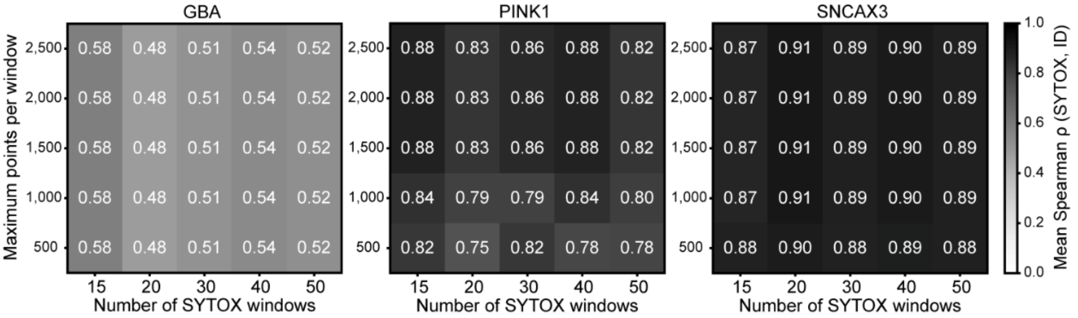


**Figure S1. Robustness of the SYTOX–ID association to window number and sample size.** Spearman correlations between SYTOX burden and MLE-based ID were evaluated across different numbers of overlapping SYTOX windows and maximum points per window. MLE was calculated with k = 20 using 10% windows after 10-NN density trimming. Values indicate mean Spearman ρ across 10 subsampling repeats.

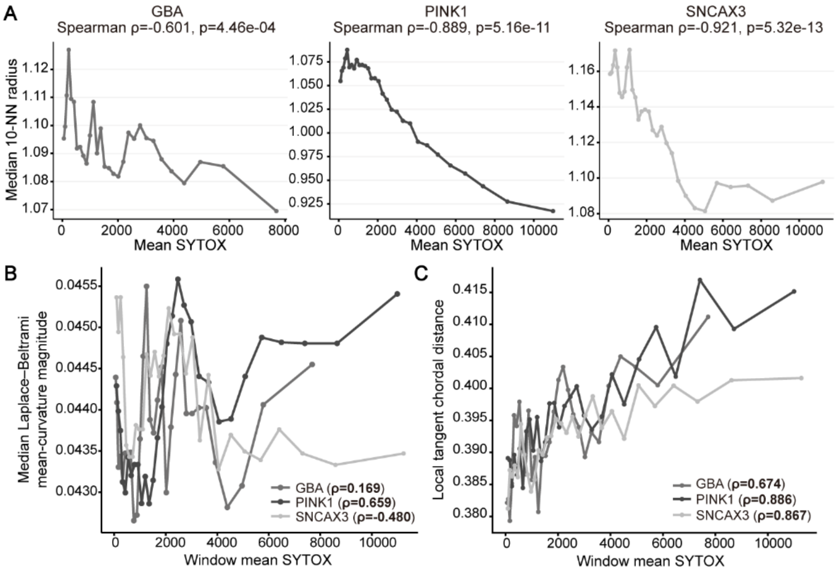


**Figure S2. Cell death burden variation in the latent manifold.** a, Changes in median 10 nearest neighbor (10-NN) radius with SYTOX burden in each subtype. Each point represents the median 10-NN radius calculated within a SYTOX ranked window. The Spearman correlation coefficient (ρ) and corresponding p-value are shown in each panel. **b,** Changes in median Laplace–Beltrami mean curvature magnitude with SYTOX burden. Each point represents the curvature magnitude calculated across SYTOX windows for each subtype and showed no significant correlation with SYTOX burden. Spearman correlation coefficient (ρ) was shown in parentheses. **c,** Changes in local tangent chordal distance with SYTOX burden, and Spearman correlation coefficient (ρ) was shown in parentheses.

### Supplementary Methods

#### Intrinsic dimensionality analysis

Changes in the intrinsic dimensionality (ID) of latent representations with cell-death burden were evaluated separately for each PD subtype (GBA, PINK1, and SNCA×3). Samples with positive intensity_per_pixel_ mean_3std values were ranked by SYTOX burden and divided into 30 overlapping windows, each spanning 10% of subtype-specific samples. Within each window, points above the 95th percentile of the 10-nearest-neighbour radius were excluded to reduce the influence of sparse outliers, and up to 2,500 points were used for ID estimation. Associations between window-level mean SYTOX burden and ID were assessed using Spearman's rank correlation coefficient (ρ). Analyses were repeated across encoder architectures and random seeds to assess reproducibility.

The Levina–Bickel maximum likelihood estimator (MLE) was used as the primary ID estimator. If $T_j(x_i)$ denotes the distance from sample $x_i$ to its $j$-th nearest neighbour, the local intrinsic dimensionality was estimated as follows[1]:

$$\hat{\mathbf{d}}_i = \frac{k-2}{\sum_{j=1}^{k-1} \log\left(\frac{T_k(x_i)}{T_j(x_i)}\right)}$$

Window-level ID was calculated as the mean of valid local estimates. The primary analysis used $k = 20$ and the analysis was repeated across multiple parameter settings to assess robustness.

TWO-NN and GRIDE were used as complementary estimators. For TWO-NN, the first- and second nearest neighbour distances for each point were denoted as $r_1$ and $r_2$, respectively, and the distance ratio $\mu = r_2/r_1$ was calculated. The final TWO-NN estimate was obtained based on the nearest neighbour distance ratio framework, in which $\mu$ follows a Pareto-type distribution, as follows[2,3]:

$$\boldsymbol{\mu}_i = \frac{r_{2,i}}{r_{1,i}}, \qquad \hat{\mathbf{d}}_{\mathrm{TWO-NN}} = \frac{n-1}{\sum_{i=1}^{n} \log\, \mu_i}$$

Here, $n$ denotes the number of valid distance ratios. Because TWO-NN uses only the two nearest neighbours, it reflects dimensionality at a highly local geometric scale.

For GRIDE, let $r_{n_1}$ and $r_{n_2}$ denote the distances to the $n_1$-th and $n_2$-th nearest neighbours, respectively, and define the distance ratio $\mu = r_{n_2}/r_{n_1}$. The dimension $d$ was then estimated numerically as the value that best fits the following likelihood. In this analysis, $n_1 = 10$, $n_2 = 20$ were used[3].

$$\boldsymbol{\mu}_i = \frac{r_{n_2,i}}{r_{n_1,i}}, \qquad \mathbf{f}(\mu \mid d) \propto d\, \frac{(\mu^d - 1)^{n_2 - n_1 - 1}}{\mu^{(n_2-1)d+1}}$$

Separately from neighbour distance based ID estimation, Shannon effective rank (eRank) was calculated to evaluate how broadly the latent variation was distributed across feature directions. The latent vectors within each window were mean centered, and covariance eigenvalues $\lambda_j$ were obtained using singular value decomposition. Shannon effective rank was then defined as follows[4]:

$$\mathbf{p}_j = \frac{\lambda_j}{\sum_k \lambda_k}, \quad \mathbf{r}_{\mathrm{eff}} = \exp\left(-\sum_j p_j \log\, p_j\right)$$

#### Exploratory analysis of latent space geometry

To determine whether increasing ID was accompanied by changes in local latent-space organisation rather than simple spatial expansion, local neighbourhood scale, Laplace–Beltrami (LB) mean-curvature magnitude, and tangent-space orientation were examined in relation to SYTOX burden.

Local neighbourhood scale was defined as the Euclidean distance from each point to its 10th nearest neighbour. The median 10-NN radius within each SYTOX window was compared with the corresponding mean SYTOX burden.

For curvature analysis, a k-nearest-neighbour graph with an adaptive Gaussian kernel was constructed from raw latent representations, followed by Coifman–Lafon normalisation to reduce sampling-density effects[5]. The LB coordinate vector was estimated as

$$\mathbf{L}(x_i) = \frac{(PX)_i - x_i}{\epsilon\sigma_i^2}, \qquad \mathbf{H}_i = \frac{||\mathbf{L}(x_i)||_2}{d_i}$$

where $X$ denotes the matrix of latent coordinates, $P$ is the density normalized diffusion transition matrix, $\epsilon$ is the kernel scale parameter, and $\sigma_i$ is the local bandwidth. Mean-curvature magnitude was estimated from the norm of the LB coordinate vector normalised by the local intrinsic dimensionality $d_i$.

Finally, local tangent-space orientation was assessed after reducing raw latent representations to 50 dimensions by global PCA[31]. Local tangent subspaces were estimated from the 30 nearest neighbours using the leading 10 local principal components. Differences between each tangent space and those of its five neighbouring tangent spaces were quantified using the principal-angle-based chordal distance[6]:

$$d_{chordal}(U,V) = \sqrt{\frac{1}{d}\sum_{j=1}^{d} \sin^2 \theta_j}$$

where $d$ is the tangent-subspace dimension and $\theta_j$ are the principal angles between tangent subspaces $U$ and $V$. Tangent-space variation was summarised within each SYTOX window and related to mean SYTOX burden using Spearman's ρ.
